# Peripheral microbial metabolites as indicators of gut microbiome disruption: systematic review and meta-analysis

**DOI:** 10.64898/2026.03.31.715561

**Authors:** Taylor Kain, Eric Armstrong, Bryan Coburn

## Abstract

**Background:** Gut microbiome disruption is often characterized by loss of obligate anaerobic bacteria, which may lead to altered production of microbial metabolites that can be detected peripherally. The application of widely used sequencing-based microbiome analyses to clinical settings is limited by cost, turnaround time, and challenges with patients with very low stool output. Since some products of strictly bacterial metabolism detectable in blood, peripheral metabolites may provide a potentially rapid and scalable indicator of gut microbiome composition and function. We performed a systematic review and meta-analysis of studies reporting circulating microbial metabolites and gut microbiome composition to evaluate whether peripheral microbial metabolites could identify gut microbiome perturbation.

**Results:** Candidate metabolites were identified systematically across an independent set of studies reporting metabolite-microbiome associations, enabling assessment of reproducibility across disease states and cohorts. We performed a meta-analysis of 19 human cohorts comprising 3242 participants with paired blood metabolite and stool microbiome data. Anaerobe depletion (obligate anaerobe relative abundance <0.70 by sequencing) was associated with decreased products of anaerobic microbial metabolism. Combinations of metabolites distinguished individuals with anaerobe-depleted microbiomes from those without. Circulating metabolite levels distinguished between cases and controls with similar performance as gut microbiome composition across a range of health/disease states, and changed markedly within patients experiencing gut anaerobe depletion after antibiotic exposure.

**Conclusions:** Circulating microbial metabolites are potentially informative indicators of gut microbiome disruption and may serve as a rapid and method for patient stratification in clinical trials or acute care settings.

**Importance:** Circulating microbial metabolites represent a practical and scalable approach to detecting significant gut microbiome disruption, particularly loss of obligate anaerobes. Unlike stool-based sequencing, which can be logistically challenging and slow, blood-based metabolite profiling could be actionably integrated into existing clinical workflows. Our findings suggest metabolites capture compositional consequences of microbiome collapse, with performance comparable to direct microbiome profiling in distinguishing disease states. Enabling diagnostic enrichment and real-time monitoring of microbiome injury (*e.g.*, during antibiotic use or critical illness) has potential implications for both clinical care and research, including selection of patients for investigation of microbiome-targeted therapies. With further validation, circulating metabolites could provide an accessible surrogate for gut microbiome composition in settings where sequencing is impractical.

## BACKGROUND

The gut microbiome plays an important role in host physiology, influencing immune homeostasis, metabolism, and resistance to pathogen colonization.^1–3^ The microbiome contributes to disease pathogenesis across a range of disease states, including inflammatory bowel disease,^4^ *Clostridioides difficile* infection,^5^ type 2 diabetes,^6^ allogeneic stem cell transplant,^7–9^ and others.^10–12^

The primary method of analyzing the microbiome in humans is either 16S rRNA gene sequencing or shotgun metagenomics.^13^ Even with recent advances in processing times, sequencing remains slower, more costly, and difficult to implement in clinically actionable time frames than conventional laboratory testing.^14^ This has impeded the translation of microbiome analysis into acute care settings, such as in acute and critical illness, where the microbiome is often highly disrupted and potentially implicated in disease risk, severity, and treatment responsiveness.^15–18^

In health, the gut microbiome is composed primarily of obligate anaerobes,^2,19,20^ and while there is no standard definition of a “healthy” microbiome,^2,21^ microbiome disruption is consistently associated with their loss, which is implicated in altered immune system signalling, susceptibility to pathogen overgrowth, and worse clinical outcomes.^7,8,17^ Microorganisms produce an array of microbial metabolites that enter systemic circulation. These metabolites have important local effects in the gut lumen,^22–24^ and also affect distal organ function and physiology.^25,26^ Unlike sequencing-based microbiome analysis, measurement of analytes in plasma or serum is routinely integrated into clinical practice, with rapid, scalable and inexpensive workflows in wide use.

We hypothesized that circulating microbial metabolite levels would correlate with gut microbiome disruption and identify individuals with substantial loss of gut obligate anaerobes. We performed a systematic review and meta-analysis to evaluate the consistency of these associations across diverse cohorts and disease states.

## METHODS

### Literature Search

To identify eligible studies, we developed a protocol in accordance with the Preferred Reporting Items for Systematic Reviews and Meta-Analyses (PRISMA) checklist,^27^ which was registered with the International Prospective Register of Systematic Reviews (reference no. CRD42024504053). We searched for relevant studies in 3 databases (MEDLINE, Embase, and Cochrane) in February 2024 (full search strategy in supplementary Table S1). We included only English language texts and excluded papers prior to the year 2000. We imported all abstracts into Covidence (Veritas Health Innovation)^28^ for review. We included studies that either directly compared circulating (*i.e.*, serum or plasma) metabolite concentrations with gut microbiome composition, or studies where both peripheral metabolite and gut microbiome data were publicly and easily available to be secondarily analyzed. Any studies that required a special access request or primary analysis of raw mass spectroscopy data were excluded. We excluded all preclinical studies as well as any studies that utilized only non-peripheral metabolite levels, or non-intestinal microbiomes. Two authors (TK and EA) independently and in duplicate screened abstracts and then full texts. We constructed a PRISMA diagram of included studies (Figure 1); studies in Box A were used to generate a list of candidate metabolites, while studies in Box B were used in the primary analysis.

**Figure 1.**
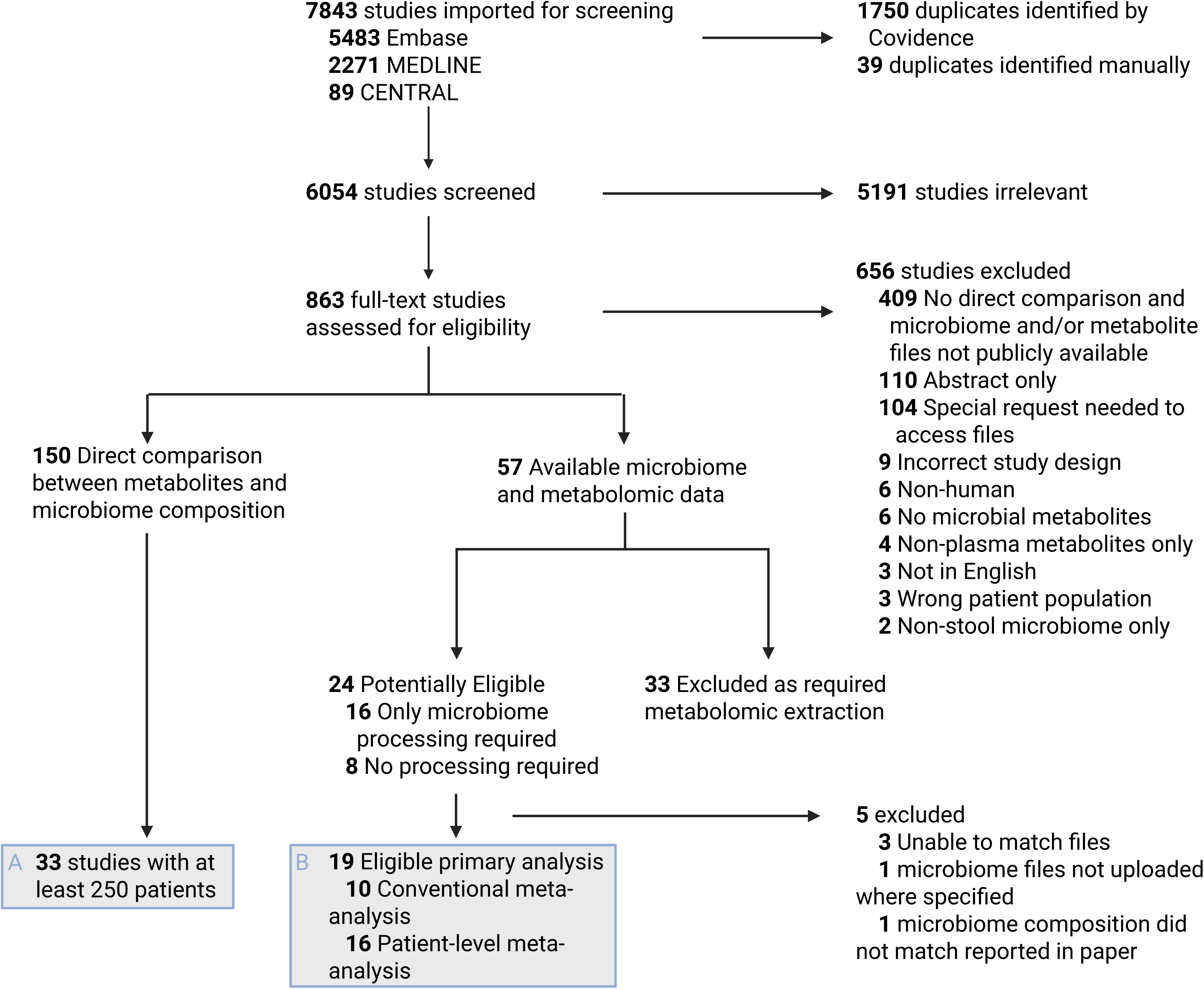
PRISMA diagram of studies screened and included in systematic review. Box A represents studies included in primary analysis, while Box B were independent studies used to generate a list of candidate metabolites used in the meta-analyses.

150 studies met inclusion criteria but did not have publicly available metabolite or gut microbiome data and instead made direct comparisons between metabolite and gut microbiome features (Box A). Details of metabolite selection are described in supplementary methods. From this analysis, 22 candidate microbial metabolites were selected, and we hypothesized that these metabolites were the most likely to correlate with gut microbiome composition in the meta-analysis. Details of the metabolites and classification are shown in supplementary Table S3. Metabolites with modifications from human metabolism (*i.e.,* addition of a sulfate group by the liver) were grouped together if only one metabolite was reported within a study.

### Metadata Extraction

For each included study, 2 reviewers (TK and EA) concurrently extracted metadata including publication year, location of study, main disease state(s) of study participants, dates of subject enrollment, study design, baseline characteristics (*e.g.*, age, sex, race, etc. where available), study interventions, and relevant clinical outcomes (if applicable). Quality assessment and risk of bias tools were not used as analysis was done with primary data.

### Microbiome Data Extraction

16S rRNA and metagenomic sequencing data were downloaded in FASTQ format from public repositories and were processed with quality filtering and denoising of reads performed. Adapters were trimmed and human reads were removed for metagenomic sequencing. Amplicon sequence variants (ASVs) were taxonomically annotated and collapsed to the genus level.^29^ Samples were rarefied to 5000 reads (16s) and 500,000 reads (metagenomics), and all samples below thresholds were excluded. Bacterial genera were classified as anaerobes, non-anaerobes, or unclassified based on descriptions in Bergey’s Manual of Systematic Bacteriology^30^ and/or supporting literature.

### Metabolite Data Extraction

Metabolite levels were extracted from public data repositories or the study’s supplementary material. The majority of studies reported metabolite levels as batch normalized and unitless values. Lower limit of detection cut-offs for each metabolite was therefore not reported. If metabolite values were reported as below the lower limit of detection, then they were imputed as half the lowest metabolite value reported for that metabolite in that study.^31,32^ To manage unitless values on different scales, we employed two techniques: 1) conventional study-level meta-analysis comparing relationships between gut microbiome composition and metabolite levels, and 2) patient-level meta-analysis after normalization with a reference metabolite.

### Data Visualization and Conventional Meta-Analysis Methods

Several studies had longitudinal samples collected. To decrease repeated measures bias, only one timepoint per patient was included other than in the longitudinal analysis. The middle time point was selected for each of these studies to increase the inclusion of more disrupted gut microbiomes in the analysis.

To predict gut microbiome disruption, an anaerobe relative abundance (RA) cutoff of 0.70 was used, approximating the lowest 10% of healthy individuals based on published data, ^2,33^ and samples were classified as “Normal/High” or “Low” based on this cut off. Further details in supplemental methods.

To examine the relationship between individual metabolites and anaerobe RA, metabolite values were normalized within each study to account for inter-study differences. Values were z-score normalized within each study, and mean z-scores were calculated per study and metabolite. A subset of metabolites: deoxycholate (DCA), ursodeoxycholate (UDCA), indole-3-propionate (I3P), hippurate, phenylalanine, p-cresol sulfate (PCS), p-cresol glucuronide (PCG), and phenylacetylglutamine (PAG) – were analyzed across studies with sufficient representation. For each study, the proportion of samples in each quadrant of anaerobe RA (high/low) versus metabolite level (above/below mean) was calculated.

### Patient-Level Meta-Analysis Methods

To combine patient-level data, we normalized metabolites using a reference metabolite level. Similar strategies have been utilized elsewhere in the literature.^37^ Further details in supplementary methods.

To evaluate the predictive performance of individual and combined metabolites for low anaerobe RA, we computed area under the curve (AUC) values for all single metabolites and all possible 2, 3, and 4 metabolite combinations. For each combination, AUCs were calculated across available samples, and distributions were visualized as boxplots. The best-performing models were selected based on Bayesian Information Criterion, which penalizes based on model complexity. Receiver operating characteristic (ROC) curves were generated for the top models using logistic regression with class weighting to account for imbalanced outcomes.

Point estimates for AUC, sensitivity, specificity, accuracy, positive predictive value, and negative predictive value were calculated on the full dataset. ROC plots display stepwise curves with shaded ribbons representing bootstrap-derived 95% confidence intervals. Two metabolite combinations were used to balance interpretability and statistical power. Metabolites were grouped by biochemical class (secondary bile acids, primary bile acids, aromatic amino acid microbial metabolites, other microbial metabolites, amino acids, and steroids) to visualize trends in predictive ability across metabolite categories. Results are presented as boxplots with overlaid points to show the distribution of AUCs for each possible metabolite combination.

### Case-Control Methods

All studies from the conventional meta-analysis that reported disease cases and controls were selected for further analysis if data was available for more than one microbial metabolite. Random-forest classification models were used to classify cases from controls using 1) gut microbiome data in the form of genus-level RA and 2) levels of all candidate metabolites. Prior to generation of random forest models, datasets were split 70/30 into test/train sets. Random forest models were generated with 10-fold cross-validation. AUC for each model was obtained. Clustering of gut microbiome data was done with principal coordinate analysis (PCoA) using Bray-Curtis dissimilarities generated from genus-level gut microbiome data. Clustering of metabolite data was done with principal component analysis (PCA).

### Longitudinal Analysis Methods

For Rashidi *et al.* metabolite abundances were log10(x+1) transformed due to skew. Patients were categorized as “Dominated” or “Never Dominated” based on *Enterococcus* domination at any time point during study. Domination was defined as *Enterococcus* RA > 30% and at least double the next most abundant taxa.

For Tanes *et al.* RA of anaerobic bacteria, Shannon diversity, and reported metabolite concentrations (indole-3-propionate, imidazole propionate, phenylalanine) were binned in 5-day intervals (0–5, 6–9, 10–15 days) relative to study initiation. Longitudinal trends were visualized with thicker LOESS smooth curves with 95% confidence intervals.

### Statistical Analysis

Statistical analysis was performed in R v4.3.1. Metabolite associations were calculated using Spearman correlations. For the conventional meta-analysis point estimates were calculated as the ratio of the median in the high anaerobe RA group over the median in the low anaerobe RA group, with a cut off of 0.70 for anaerobe RA. Similar strategies have been utilized elsewhere, with some modelling data showing that median strategies may be preferrable to transformation strategies,^38–40^ when outcome data is skewed.^39^ Confidence intervals were calculated using a nonparametric bootstrap (1,000 replicates). For pooled estimates of individual metabolites, an inverse-variance weighted fixed-effects model was used. Ratios were visualized on the log scale, but estimates are reported untransformed. Details are shown in the results section and the legends of Figure 2 as well as supplementary Figure S2. To visualize relationship of metabolites, samples were z-score normalized. L-shaped associations were evaluated using Fisher’s exact test to estimate odds ratios. Details shown in legend of Figure 3.

**Figure 2.**
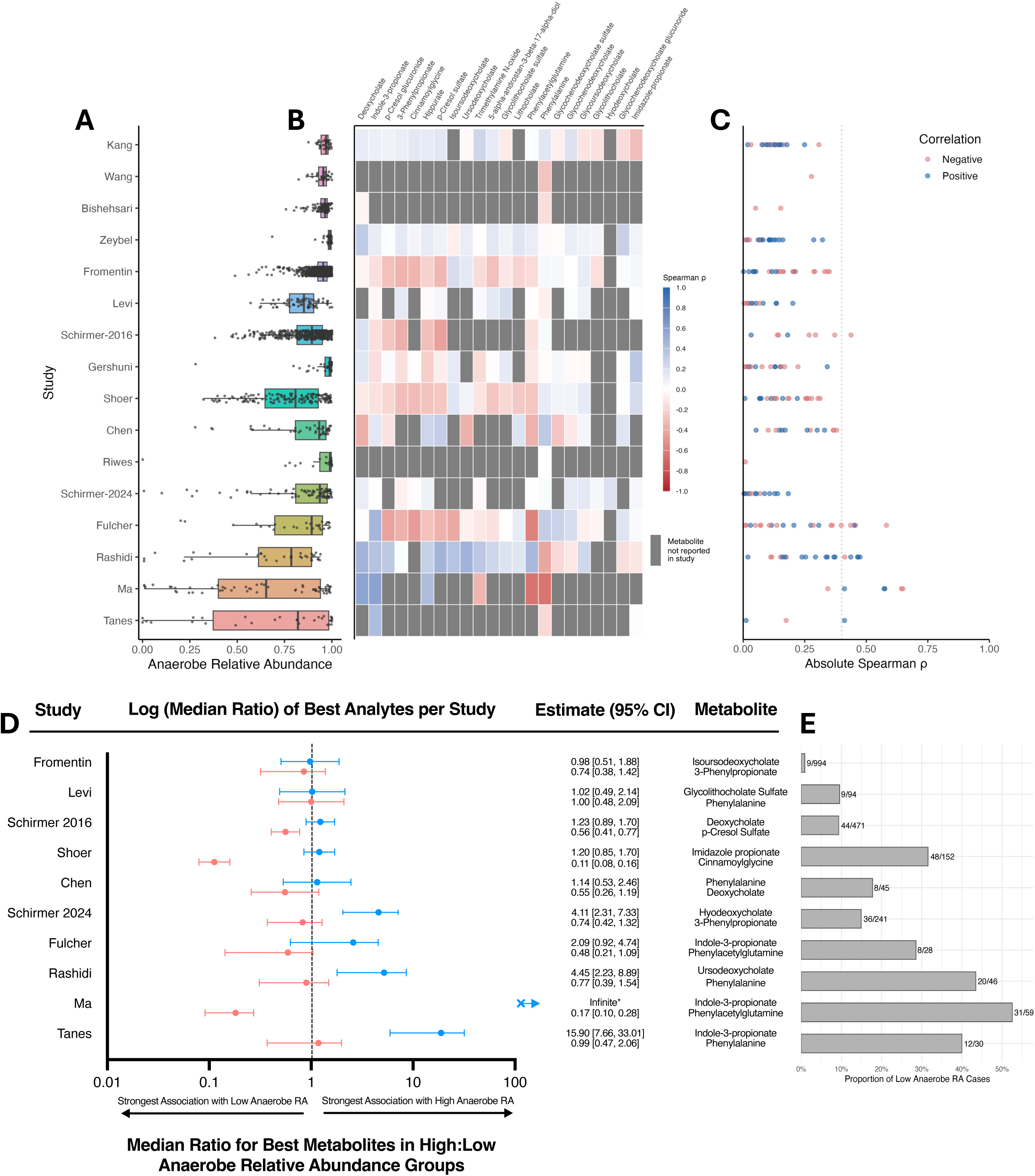
Correlations of metabolites and anaerobe relative abundance (RA) and conventional meta-analysis of the best-performing analytes. A) Anaerobe RA by study, in order of increasing variance. B) A heatmap showing Spearman correlations between each metabolite and anaerobe RA. Grey boxes indicate that the metabolite was not reported in that study. C) Absolute Spearman correlations between peripheral metabolites and stool anaerobe RA. Dotted line shows Spearman > 0.4. D) Forest plot showing the median ratio between anaerobe RA > 0.70 or < 0.70 most positively and negatively correlated metabolites in each study. Confidence intervals were calculated using bootstrapping. E) Proportion of participants with anaerobe RA < 0.70 in each study. *Indole-3-propionate in Ma *et al.* had an infinite median ratio as all patients in the low anaerobe RA group had a level of 0.

**Figure 3.**
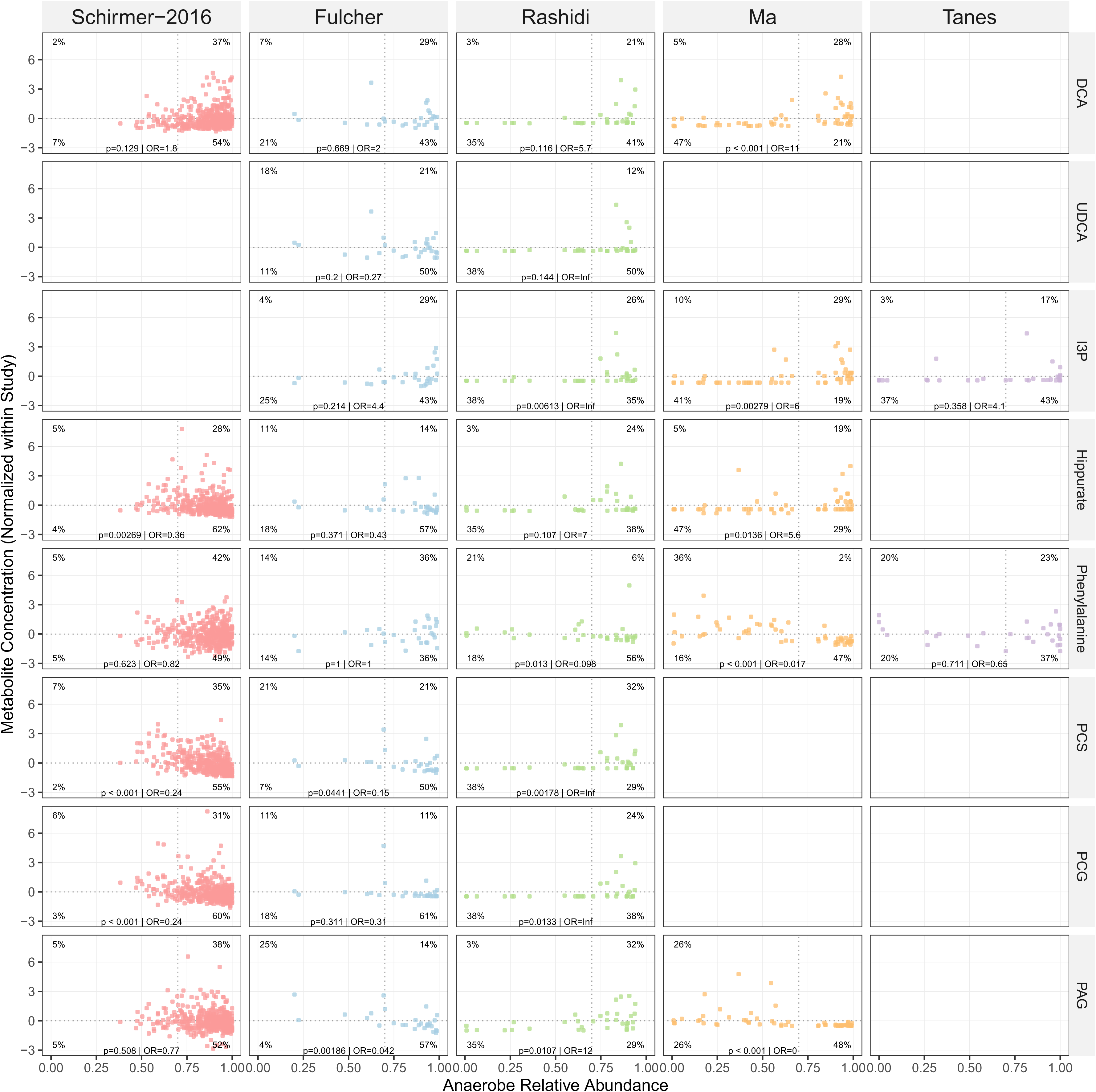
Normalized metabolite concentrations and anaerobe relative abundance (RA). Relationships between metabolite levels correlated with anaerobe RA (included studies had minimum Spearman >0.40 in at least one metabolite). Quadrants are split based on median metabolite level (in each study, y-axis) and anaerobe RA of 0.70 (x-axis), with percentage of samples shown in each quadrant. P-values are from Fisher’s exact tests. DCA = deoxycholate, UDCA = ursodeoxycholate, I3P = indole-3-propionate, PCS = p-cresol sulfate, PCG = p-cresol glucuronide, PAG = phenylacetylglutamine

For patient-level meta-analysis, results were normalized based on the reference metabolite, histidine. Performance of ROC models were quantified by AUC and McFadden pseudo-R^2^. To account for increasing complexity of models, we computed adjusted pseudo-R^2^ and Bayesian Information Criterion (BIC). Combinations were ranked primarily by BIC, penalizing larger more complex models. Predictions were assessed on out-of-bag samples from 500 bootstrap resamples to compute 95% confidence intervals for sensitivity and specificity at each threshold. Details can be found in results section and legend for Figure 4.

**Figure 4.**
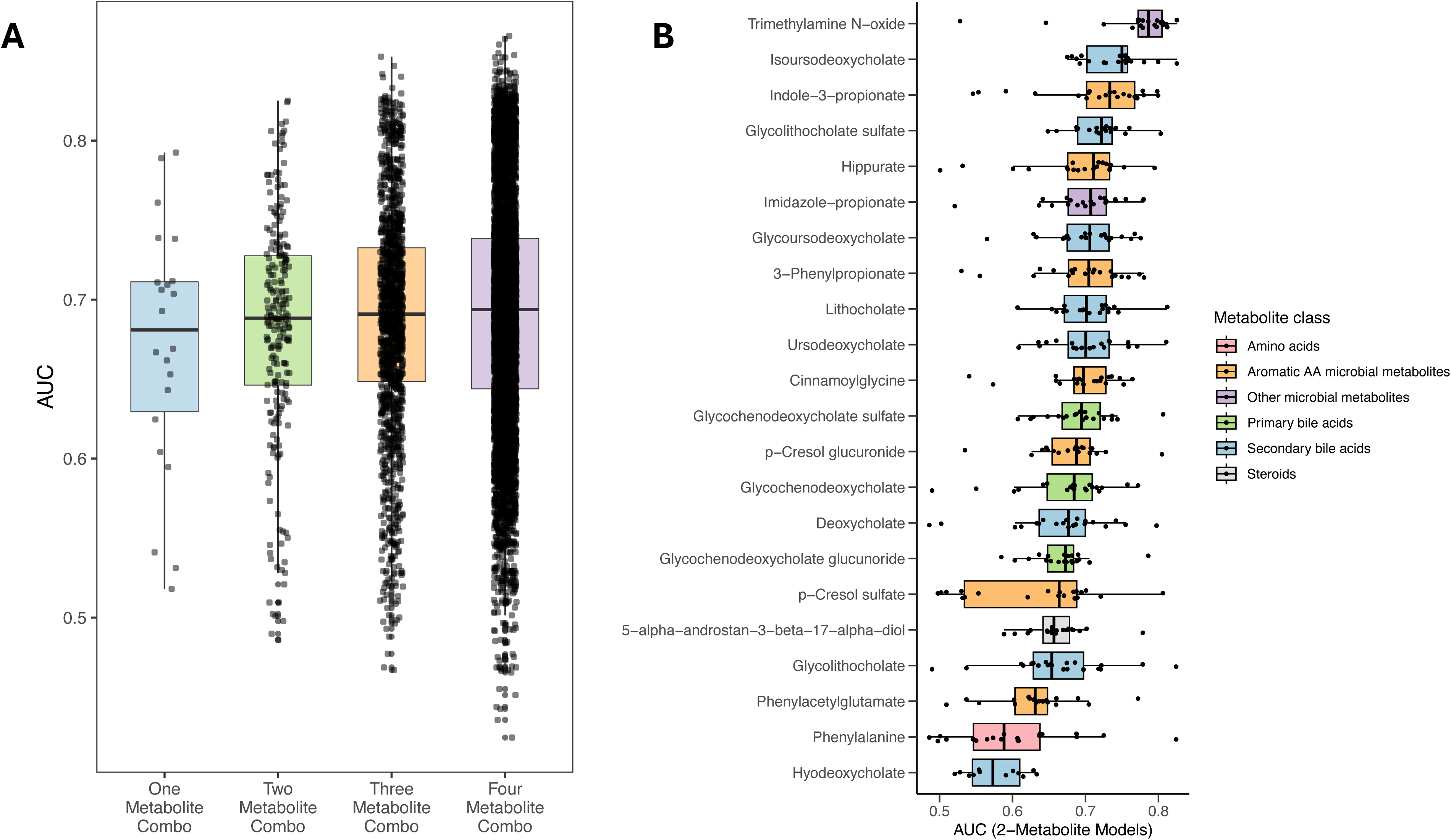
Prediction of low anaerobe relative abundance (RA) using microbial metabolites. A) areas under ROC curves (AUCs) for individual metabolites and all metabolite combinations to predict anaerobe RA <0.70. B) Boxplots showing distribution of AUCs for all two-metabolite combinations involving the reference metabolite. Metabolites are ordered by median AUC. Metabolites are coloured by metabolite class. AA=amino acid, AUC=area under the curve.

For case-control analysis, differentially abundant bacterial genera were measured between cases and controls using Multivariable Association with Linear Models 2 (MaAsLin2), with case-control status as fixed effect and a false discovery rate (FDR)-adjusted *q* value threshold of 0.10 for significance testing. Metabolite abundances were compared between cases and controls with Mann-Whitney U tests and *p* values were adjusted for multiple comparisons with the FDR. For metabolite comparisons, an FDR-adjusted *q* value threshold of 0.05 was used for significance testing. Details can be found in results section, as well as legends for Figure 5, and supplementary Figure S5.

**Figure 5.**
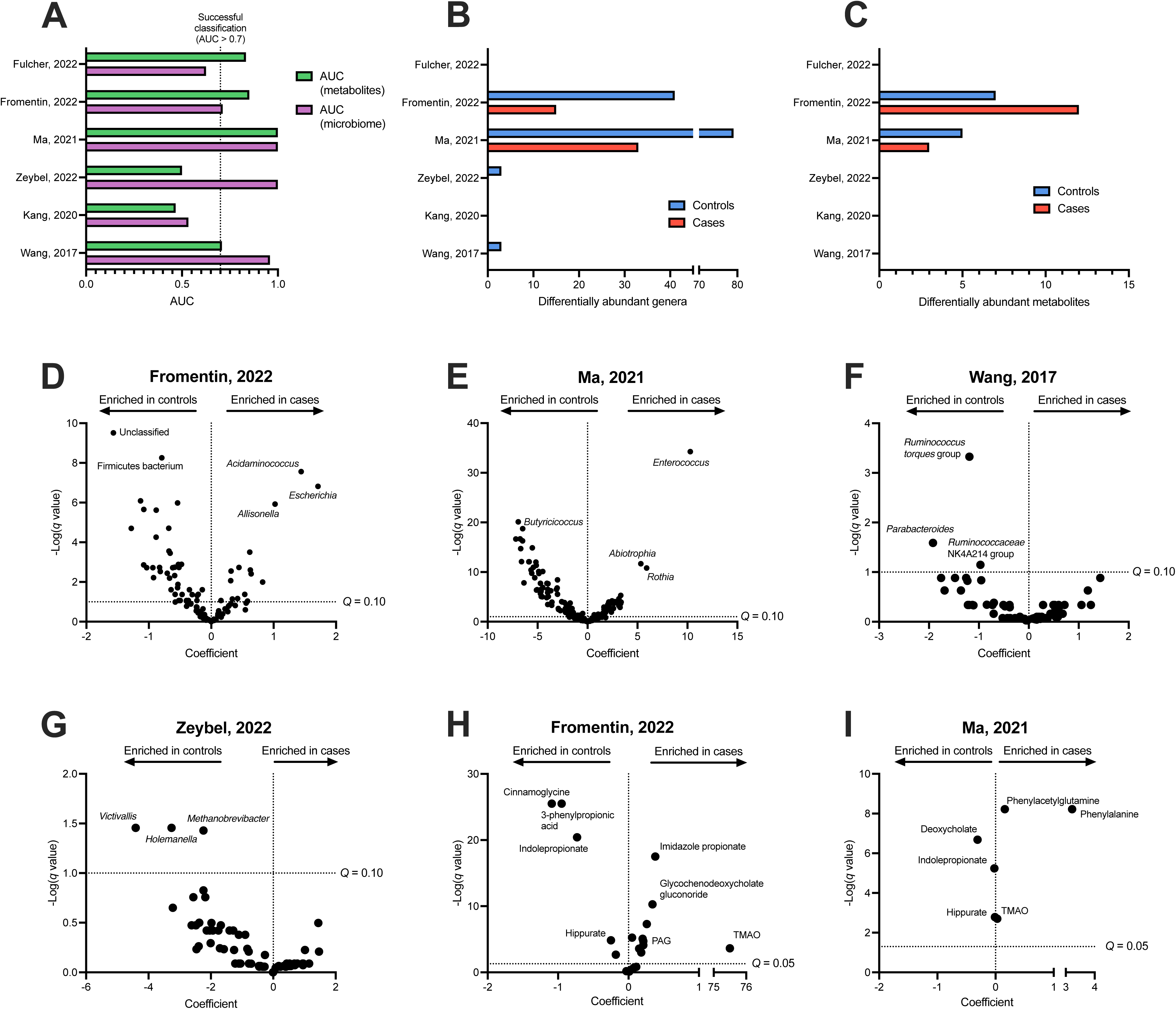
Differentiation of disease cases from controls using the gut microbiome and peripheral microbial metabolites. A) AUCs of gut microbiome and metabolite random forest classifiers for cases and controls, with the threshold for successful classification at AUC = 0.7. Number of differentially abundant B) bacterial genera and C) microbial metabolites in cases and controls. D-G) Association between individual bacterial genera and case-control status as determined with MaAsLin2, with an FDR-corrected q value of 0.10 used as the threshold for significance testing. H-I) Association between individual candidate metabolites and case-control status as determined with Mann-Whitney U tests, with an FDR-corrected q value of 0.05 used as the threshold for significance testing. TMAO=trimethylamine-N-oxide, PAG=phenylacetylglutamine.

For longitudinal analysis in Rashidi *et al.* linear mixed-effects models evaluated temporal trajectories, including fixed effects for patient group, study day, and their interaction, and a random intercept per patient. The interaction term tested for group differences in longitudinal trends. For Tanes *et al.* pairwise comparisons between consecutive bins were conducted using Wilcoxon rank-sum tests, with significance annotated on plots. Details can be found in results section and legend for Figure 6 as well as supplementary Figure S6 and S7. Unless otherwise stated, statistical significance was defined as two-sided α < 0.05.

**Figure 6.**
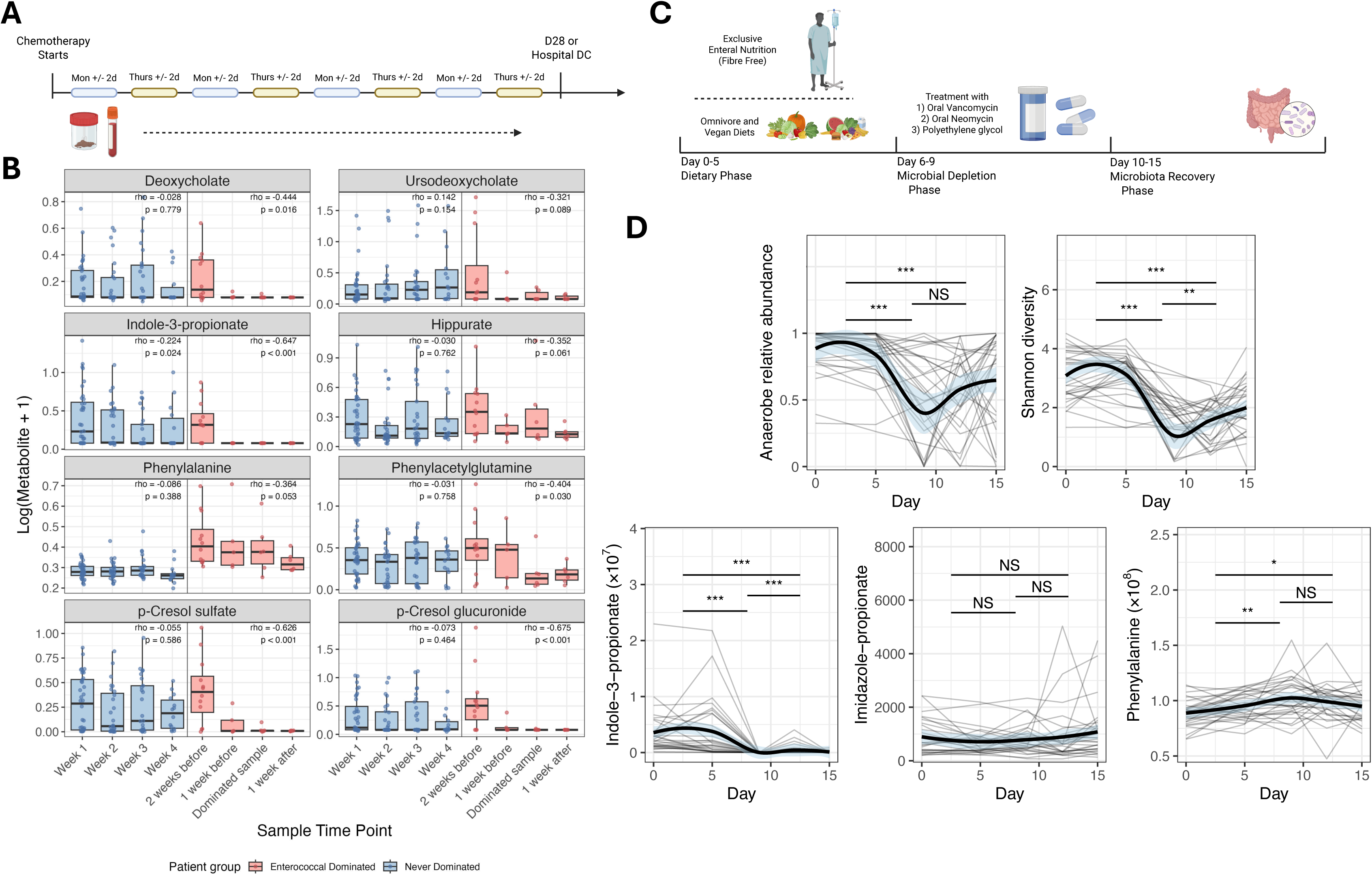
Longitudinal microbial metabolite dynamics. (A) Rashidi *et al*. study design: patients with acute myeloid leukemia (AML) were enrolled at the start of induction chemotherapy, with blood and stool collected twice weekly until hospital discharge or day 28. (B) Metabolite abundances stratified by *Enterococcus* domination status in Rashidi *et al*. Boxplots show log₁₀(metabolite + 1) across time bins, aligned to the first dominated sample for dominated patients; points denote individual samples. Spearman ρ and p-values summarize within-group temporal trends. The eight metabolites shown had |ρ| > 0.4 with anaerobe relative abundance in at least one study. (C) Tanes *et al*. study design: Participants were randomized to a dietary intervention (days 0–5), then all were given antibiotics (days 6–8), followed by gut microbiome sampling (days 10–15). (D) Longitudinal trajectories of ecological metrics and metabolite levels are shown. Thin lines show individual subject-level measurements with thick lines show locally estimated scatterplot smoothing and 95% CIs. Pairwise differences between bins (0–5, 6–9, 10–15) were tested by Wilcoxon rank-sum (p > 0.05 NS [not significant], p < 0.05 *, <0.01 **, <0.001 ***).

### Software

FASTQ files were processed with Quantitative Insights Into Microbial Ecology 2 (QIIME 2, v2024.2) using default parameters.^41^ Quality filtering and denoising of reads was performed with DADA2.^42^ ASVs were taxonomically annotated using the SILVA 132 99% database.^29^ Adapters were trimmed using Trimmomatic version 0.39^43^ and human reads were removed with KneadData version 0.12.0. Taxonomic annotation was performed with Metagenomic Phylogenetic Analysis (MetaPhlAn) version 4.1.1.^44^

Analyses were performed in R v4.3.1 using dplyr 1.1.4,^45^ ggplot2 3.5.1,^46^ pROC 1.18.5,^47^ patchwork 1.3.1,^48^ ggnewscale 0.5.2,^49^ purrr 1.0.2,^50^ caret 7.0-1,^51^ randomForest 4.17-1.2,^52^ nlme 3.1-166,^53^ ecodist 2.1.3,^54^ vegan 2.6-10,^55^ FactoMineR 2.11^56^, missMDA 1.19,^57^ and MaAsLin2 1.16.0.^58^

## RESULTS

### Systematic Review

6054 unique abstracts were screened in duplicate. After screening, 5191 studies were deemed irrelevant and 859 studies were assessed by full text for eligibility. The full search strategy is shown in supplementary Table S1. Of these studies, 150 presented only direct comparisons between metabolites and microbiome composition in the text, of which 33 studies included at least 250 participants for a total of 32607 participants. The most common populations studied were cardiovascular or metabolic diseases (12 studies, 13639 participants), with healthy cohorts being the next most common (10 studies, 9402 participants). Other populations included individuals with infection, rheumatologic, endocrine, renal, neurologic, and liver diseases. Further details of studies and articles are available in supplementary material (Box A Studies supplemental file). We used these studies to identify candidate microbial metabolites by assessing reported associations between circulating metabolites and microbiome features (additional details in Methods), with the goal of reducing the number of candidate metabolites and enriching for those most likely to be generalizable. This analysis yielded 22 candidate metabolites that were selected based on their correlations with microbiome composition (Box A; Figure 1). The selection scoring tool is shown in Table S2 and classification of candidate metabolites in Table S3. Candidate metabolites derived from this analysis were included in a formal meta-analysis of studies reporting both metabolite concentrations in plasma or serum and stool microbiome composition.

Microbiome and metabolomic data were available at an individual patient level for 57 studies. 33 of these studies required primary metabolomic data extraction and were excluded. The remaining 24 studies were reviewed for inclusion in the primary analysis. Of the 24 studies reviewed, 5 studies were excluded: 3 did not provide metadata to match metabolite and microbiome data for individual participants;^59–61^ one did not provide microbiome data in the referenced database,^62^ and; one provided microbiome data that, upon analysis, was markedly different than the published results and not consistent with a human gut microbiome.^63^ The authors for all 5 of these studies were contacted and only one responded^60^ for which an external application was required to access matching patient level details with use restrictions and therefore the study was excluded. This left 19 eligible studies^34–36,64–79^ included in the primary analysis (Box B). These 19 studies included 3242 participants, with 2999 complete cases (values present for both metabolites and microbiome for same sampling timepoint). Of these 19 studies, three did not report any candidate metabolites from our metabolite screen (selected from Box A in Figure 1) and were therefore not included in subsequent analyses.

Disrupted microbiome cases were defined as those with an anaerobe relative abundance (RA) below 0.70, which approximates the lowest 10% of the distribution of anaerobe RA for healthy individuals based on published data (supplementary methods). Six studies had fewer than 5 cases using this definition of low anaerobe RA and were therefore excluded. In total, 10 studies representing 2270 participants and 1903 complete cases were included in the study-level meta-analysis, while 16 studies, representing 2622 patients (2194 complete cases), were included in the patient-level meta-analysis. Details of the included studies for each analysis are shown in Table S4.

### Study characteristics

Characteristics of the studies included in the study-level meta-analysis are shown in Table 1. Most studies were from the United States (8/19, 41%) or China (5/19, 26.3%). Healthy populations and a variety of disease states were captured, with cardiovascular disease and cancer being the most common (both 3/19, 15.8%). Other disease states included infection, neurological, endocrine, kidney, liver, and gastrointestinal diseases. Only 2 studies (Rashidi *et al.^79^* and Tanes *et al.^36^*) reported participant-level antibiotic use before or during the study period.

**Table 1.** Overview of included studies. Age is median in years (IQR); a cell is recorded as NA if age or sex is not reported at the patient level for that study. N is the total number of unique patients. Complete cases are unique samples with paired metabolite and gut microbiome data. Dominance is defined as presence of a non-obligate anaerobe with RA > 30% and at least twice the next most prevalent taxon. IQR = interquartile range, CV = cardiovascular, MetS = Metabolic syndrome, NAFLD = Non-alcoholic fatty liver disease, RA = anaerobe relative abundance, Abx = antibiotics, PWH = persons with HIV, ARV = anti-retrovirals, ASCT = allogeneic stem cell transplant, AML = acute myeloid leukemia

| # | Author | N (complete) | Age (IQR) | Sex (% Male) | Abx Reported | Disruption Features in > 5 pts | Country/Region | Population | Cases | Comparison/Controls |
| --- | --- | --- | --- | --- | --- | --- | --- | --- | --- | --- |
| 1 | Wang<sup>78</sup> | 40 (35) | 36 (30-43) | 79.5% | No | No | China | Liver | Hepatitis B | Healthy Controls |
| 2 | Bishehsari<sup>77</sup> | 123 (118) | 44 (34-53) | 27.6% | No | No | USA | CV | MetS | - |
| 3 | Zeybel<sup>76</sup> | 41 (41) | 40 (33-45) | 68.3% | No | No | Turkey | Liver | NAFLD w steatosis | NAFLD without steatosis |
| 5 | Lin<sup>75</sup> | 517 (498) | 53 (51-54.5) | 0% | No | Anaerobe RA < 0.7 | China | Hormonal | Menopause | - |
| 6 | Schirmer 2016<sup>34</sup> | 488 (441) | 23 (21-27) | 43.2% | No | Anaerobe RA < 0.7 | Western Europe | Healthy | Healthy | - |
| 7 | Chen<sup>74</sup> | 45 (45) | - | - | No | Anaerobe RA < 0.7 | China | Oncology | Lung Cancer | Healthy Controls |
| 8 | Fulcher<sup>73</sup> | 28 (28) | - | 100% | No | Anaerobe RA < 0.7 | USA | Infection | HIV acquisition | Longitudinal |
| 9 | Fromentin<sup>72</sup> | 1087 (992) | 60 (52-66) | 61.4% | No | Anaerobe RA < 0.7<br>Dominance > 30% | Western Europe | CV | CV Diseases | Between CV Diseases |
| 10 | Tanes<sup>36</sup> | 30 (30) | 25.5 (24-37) | - | Yes | Anaerobe RA < 0.7<br>Dominance > 30% | USA | Healthy | Diets | Longitudinal<br>Interventional Design |
| 11 | Sereti<sup>71</sup> | 137 (130) | 62 (55-68) | 97.8% | No | No | Netherlands | Infection | PWH | Healthy Controls |
| 12 | Ma<sup>35</sup> | 59 (58) | - | - | No | Anaerobe RA < 0.7<br>Dominance > 30% | China | Renal | Renal Transplant | Healthy Controls |
| 13 | Uema<sup>70</sup> | 140 (140) | 60 (47-67) | 50% | No | Anaerobe RA < 0.7 | China | CV | MetS | Healthy Controls |
| 15 | Riwe<sup>69</sup> | 25 (25) | - | - | No | No | USA | Oncology | ASCT | Longitudinal of<br>Various Diets |
| 16 | Gershuni<sup>68</sup> | 48 (44) | 30 (26.5-34.5) | 0% | No | No | USA | Healthy | Obesity in Pregnancy | Normal BMI in pregnancy |
| 18 | Shoer<sup>67</sup> | 152 (148) | - | - | No | Anaerobe RA < 0.7 | Israel | Healthy | Diet | Long. of Various Diets |
| 19 | Rashidi<sup>79</sup> | 46 (34) | 59 (50-68.5) | 64.9% | Yes | Anaerobe RA < 0.7<br>Dominance > 30% | USA | Oncology | AML | Long. through AML<br>Induction Course |
| 20 | Schirmer 2024<sup>66</sup> | 241 (85) | 13 (10-15) | 55.6% | No | Anaerobe RA < 0.7<br>Dominance > 30% | USA | Gut | IBD | Weeks from UC<br>Diagnosis |
| 21 | Levi<sup>65</sup> | 94 (68) | 37 (30-46.5) | 31.9% | No | Anaerobe RA < 0.7 | Israel | Neuro | MS | Healthy Controls |
| 22 | Kang<sup>64</sup> | 38 (38) | 10.9 (8.8-12.7) | 86.8% | No | No | USA | Neuro | Autism | Healthy Controls |

Two studies included paediatric populations, and 3 studies had a median age over 60 years. The median age of participants in the 14 studies reporting participant-level age data was 52 (IQR 28-60). In reporting studies, and 1370/2961 (46.3%) were male (range 0% to 100%). No studies reported transgender individuals. Median peripheral metabolites reported across studies was 655 (range 5 [all bile acids] to 10,026). Candidate metabolites reported in each study are shown in Tables S4 and S5.

Anaerobe RA and Shannon diversity index (SDI) varied across studies (supplementary Figure S1A and S1B). The median SDI ranged from 1.65 to 5.18. Anaerobe RA varied with medians between 0.64 and 0.99 with minimum 0 and maximum 1 (Figure 2A). Correlation between SDI and anaerobe RA are shown in supplementary Figure S1C. Two studies (Rashidi *et al.^79^* and Ma *et al.^35^*) had median anaerobe RA less than 0.8. Twelve studies had at least 5 cases with anaerobe RA < 0.70.

### Metabolite Associations and Meta-Analysis

We reasoned that peripheral metabolite levels would differ most between preserved and highly disrupted microbiomes. Therefore, we hypothesized that associations would only be present in studies in which a sufficient proportion of participants had loss of anaerobes (and therefore studies with larger variance in anaerobe RA). Anaerobe RA for each study is presented in Figure 2A. Correlations between each metabolite of interest and anaerobe RA are shown in Figure 2B and 2C. Studies with low variance in anaerobe RA had weak correlations between metabolites and anaerobe RA. Five studies (Schirmer 2016,^34^ Fulcher,^73^ Rashidi,^79^ Ma,^35^ and Tanes^36^) had at least 1 metabolite with a moderate Spearman correlation (>|0.4|) with anaerobe RA, which included: deoxycholate (DCA), ursodeoxycholate (UDCA), indole-3-propionate (I3P), hippurate, phenylalanine, p-cresol sulfate (PCS), p-cresol glucuronide (PCG), and phenylacetylglutamine (PAG). The strongest negative and positive correlations with anaerobe RA were both from Ma *et al.^35^* in which PAG was negatively correlated (Spearman correlation = -0.65) and I3P was positively correlated (Spearman = 0.58).

Methods for metabolite quantification and normalization differed across studies. To control for the variable scales of each metabolite, we calculated the median ratio between metabolites in the low and high anaerobe RA groups, a strategy that has been used previously (Figure 2D).^38–40^ We again noted that larger ratios were generally seen in studies with more variance in anaerobe RA. Metabolites that were products of obligate anaerobe metabolism generally had larger ratios than other metabolites, such as those produced by metabolism by facultative anaerobes. The predicted direction of metabolite associations with anaerobe RA are shown in Table S6.

In study-level meta-analyses, log median ratios of two candidate metabolites were significantly different between high and low anaerobe RA participants: DCA (pooled estimate=1.62-fold, 95% CI: 1.32 – 2.04) and PCG (pooled estimate of 0.78-fold (95% CI: 0.68 – 0.89 fold, supplementary Figure S2). Other candidate metabolites ratios did not significantly differ between high and low anaerobe RA participants.

In studies with a greater proportion of patients with anaerobe-depleted microbiomes, products of anaerobic microbial metabolism (I3P, DCA, UDCA, and hippurate) were more strongly associated with higher anaerobe RA (Figure 3). Given that relationships between metabolites and anaerobe RAs were non-linear (i.e. decreased markedly at lower anaerobe RAs) we evaluated associations using an anaerobe RA threshold of 0.70. Across cohorts, associations between mean metabolite values and this threshold were strongest for DCA (odds ratio [OR] ranged from 1.8 [95% CI 0.86-4.1, p=0.13] to 11.4 [95% CI 2.6-72.7, p<0.001]), and I3P (OR ranged from 4.1 [95% CI 0.37-218.0, p=0.36] to infinite [95% CI 1.63-infinite, p=0.006]). Conversely, PCS, PCG, and PAG generally showed inverse associations with anaerobe RA, with the exception of Rashidi *et al.* ^79^. These effects were most notable for PAG (OR ranged from 0.77 [95% CI 0.38-1.55, p=0.51] to 0 [95% CI 0-0.18, p<0.001]). Effect sizes generally increased with higher variance in anaerobe RA across cohorts.

### Patient Level Meta-Analysis

We next generated logistic regression models of metabolite levels to predict low anaerobe RA (<0.70). To control for differences in scaling between studies and unitless reporting of metabolites, all metabolite levels were normalized by histidine (see Methods). Models included individual metabolites and combinations of up to four different metabolites. When penalizing for model complexity using Bayesian Information Criterion (BIC), 2-metabolite combinations provided the best balance between predictive model performance and parsimony (Figure 4A). AUC curves for prediction of low anaerobe RA using individual peripheral metabolites are shown in supplementary Figure S3. The best performing individual metabolites were isoursodeoxycholate (IUDCA, AUC 0.79, 95% CI 0.72-0.85) and trimethylamine-N-oxide (TMAO, AUC 0.79, 95% CI 0.75-0.83). Amongst all 2-variable models, models which included TMAO had the best median performance (median AUC 0.79, interquartile range [IQR] 0.77-0.81, Figure 4B), and inclusion of multiple secondary bile acids in 2-variable models was associated with better performance of those models (Figure 4B). Combinations such as phenylalanine + glycolithocholate (GLCA) and IUDCA + TMAO produced the strongest predictions (supplementary Figure 4A and 4B) among 2-metabolite combinations. Sensitivity analysis using alanine normalized metabolites levels produced similar results (examples shown in supplementary Figure S4C and S4D).

### Application of peripheral metabolite levels to diagnostic scenarios

#### Case-Control Prediction

To test whether circulating metabolites and gut microbiome composition could distinguish between health and disease, we selected all studies with data for disease cases and controls that reported at least two candidate microbial metabolites. Six studies across diverse participant populations were eligible for inclusion in this analysis (Table S5). We used random forest classification models with genus-level gut microbiome data or peripheral microbial metabolites to classify cases from controls in each study. Case and control status was successfully predicted (AUC > 0.7) in 4/6 microbiome classifiers and 4/6 metabolite classifiers (Figure 5). Random forest classification of cases and controls in testing datasets are presented in Table S7. Clustering of cases and controls based on microbiome or metabolite data followed similar patterns as the random forest classifiers; studies with successfully classified cases and controls tended to have clearer separation of cases and controls in PCoA (for microbiome data) or PCA (for metabolite data) (supplementary Figure S5). We then used MaAsLin2 (for microbiome data) or Mann-Whitney U tests (for metabolomic data) with FDR correction to test for differentially abundant features between cases and controls. Gut commensals were more abundant in controls over cases, and cases were enriched with pathobionts such as *Enterococcus* and *Escherichia* (Figure 5). Differentially abundant metabolites between cases and controls were observed in two studies, Ma *et al.^35^* and Fromentin *et al*.^72^ Although these studies measured different panels of metabolites, cases in both studies had elevated TMAO and PAG and controls had elevated I3P and hippurate (Figure 5). Peripheral metabolites performed similarly to gut microbiome for distinguishing between cases and controls in these datasets.

#### *Analysis of* longitudinal *sampling with acquired anaerobe loss*

Two studies (Rashidi *et al.^79^*and Tanes *et al.^36^*) collected samples longitudinally and included participants that developed microbiome disruption during sampling. Rashidi and colleagues reported microbiome composition and metabolites for patients being treated for newly diagnosed AML (Figure 6A). Many participants were given antibiotics during their acute illness which resulted in decreased microbiome diversity and anaerobe RA (supplementary Figure S6) and concomitant decreases in microbial metabolites produced by anaerobes. Notably, in this population, the most common taxon to emerge as the dominant one was *Enterococcus*, which has been linked to multiple poor outcomes in similar disease states.^7,8^ In patients with acquired *Enterococcus* dominance, defined as *Enterococcus* RA >0.30 and at least twice as abundant as the next most abundant taxon, most metabolite levels dropped to undetectable levels at the time of *Enterococcus* dominance or preceding this point (Figure 6B). Unexpectedly, we observed significant reductions in metabolites that were negatively associated with anaerobe RA in the meta-analysis, such as PAG, PCS, and PCG. These metabolites (or precursors to these metabolites) can be produced by facultative anaerobes when substrates accumulate with obligate anaerobe loss, a process which may be disrupted by antibiotic-mediated pathogen depletion. Importantly, all study participants in this study were given fluroquinolone prophylaxis and consequently, expansion of facultative anaerobic taxa was rare (supplementary Figure S6), potentially explaining this observation.

We next examined the relationship between metabolites and microbiomes after a dietary intervention and antibiotic exposure in healthy volunteers in the study reported by *Tanes et al.^36^*Antibiotics (vancomycin 500mg orally q6h and neomycin 1000mg orally q6h) and oral polyethylene glycol (PEG) were administered to study participants on days 6, 7, and 8 (Figure 6C). As expected, profound drops in diversity and anaerobe RA during and after administration of antibiotics and PEG was evident (Figure 6D). Concordantly, I3P also dropped to undetectable levels in all participants after antibiotic administration, while as expected the same effect was not seen for imidazole propionate or phenylalanine (Figure 6D). Individual patient metabolite dynamics are shown in supplementary Figure S7.

## DISCUSSION

In this systematic review and meta-analysis, we found that circulating microbial metabolites were associated with profound gut anaerobe loss, a finding driven by studies where a significant proportion of participants had highly disrupted microbiomes. Products of anaerobic microbial metabolism, including unconjugated secondary bile acids, indole derivatives, and hippurate, were most consistently associated with microbiome disruption. The metabolites we assessed largely fell into one of three categories: 1) products of obligate anaerobe metabolism, which decrease with anaerobe loss, 2) products of facultative anaerobe metabolism, which generally increase with anaerobe loss and pathobiont overgrowth, or 3) substrates of anaerobic microbial metabolism.

Metabolites in group 1 include secondary bile acids (SBA, such as DCA, UDCA) and indole metabolites (such as I3P), which were consistently reduced with anaerobe depletion in our study. Primary bile acids (PBA) are deconjugated by taxonomically diverse organisms, and then converted to SBA,^80,81^ a process largely restricted to obligate anaerobes within the *Clostridia* class.^82,83^ I3P is formed in the gut via reductive microbial metabolism of tryptophan, a process carried out by a restricted group of obligate anaerobes.^84^ SBA have been shown to have anti-microbial properties,^85–87^ anti-inflammatory effects,^81^ and play a key role in colonization resistance.^88^ I3P stimulates goblet cells to secrete mucin and enhance expression of tight junctions, which is linked to inflammation and immune activation.^84,89^ I3P has been shown in various animal models and human observational data to have beneficial effects on distal organs.^90–94^ The associations we observe therefore represent both potentially informative indicators of microbiome composition and host-biology-relevant function.

Metabolites from group 2 tended to increase with anaerobe loss, although this association was not as uniform across different studies. The precursor to PAG, phenylacetate (PAA) is produced primarily by obligate anaerobes, such as *Clostridia* and *Bacteroides* species before being conjugated in the liver. When obligate anaerobes are depleted, phenylalanine is rerouted to PAA via facultative anaerobes.^95^ Obligate anaerobes also normally consume PAA, so their loss leads to accumulation of PAA and thus PAG. Its production depends on other factors such as antibiotic exposure, kidney injury, and organ dysfunction, which were not reliably reported at the patient level in the studies included in our systematic review. For example, in Rashidi *et al.*,*^79^* PAG decreased with gut microbiome disruption, the opposite direction to associations in other studies. Of note, the patients within this study were all treated with fluoroquinolone antibiotics and consequently had very low RA of *Enterobacteriaceae*. The variability in this association may represent biologically informative heterogeneity, but this needs to be assessed in larger and specifically designed studies. TMAO, which can be produced via facultative anaerobe metabolism, was also among the strongest predictors of low anaerobe RA and was enriched in disease cases. Finally, metabolites from group 3, including PBAs (such as glycochenodeoxycholic acid) and essential amino acids, showed the weakest or least consistent associations. Although it is logical that substrates of anaerobic microbial metabolism would accumulate in response to gut anaerobe depletion, a variety of factors independent of microbial metabolism may affect substrate levels including diet, host production, and reabsorption from the intestinal lumen.

Our study has important limitations. First, only a minority of studies included populations with a large proportion of individuals with microbiome disruption. Therefore, the associations we present are driven by a smaller number of individuals from fewer disease states, despite the large number of individuals in the overall meta-analysis. Second, our initial candidate-selection strategy was designed to emphasize robustness (reproducibility across a large number of candidate-selection studies), potentially excluding highly informative metabolites represented in fewer published reports. Third, metabolites were largely reported as batch-normalized, unitless values, preventing a direct analysis of merged data, and while normalizing data by study would control for inter-study differences in methods, it may also eliminate important inter-study differences in analyte (and microbiome) distributions, potentially leading to false-negative associations. Fourth, we used simplified indicators of microbiome disruption with an arbitrary threshold based primarily on relative abundance distributions rather than on more detailed/nuanced risk-associated microbiome-biology or microbiome-health associations. There may exist specific and preserved relationships between metabolites and individual families or taxa not detected with such a broad ecological indicator. Finally, there was very limited information on potentially informative covariates, such as age, sex, race, and diet among studies, which may have driven within- and between-study metabolite/microbiome associations. Although our observations are therefore broadly representative, it is possible that the influence of some of these variables, such as diet or antibiotic exposure, may alter microbiome-metabolite associations in a clinically important way. These limitations indicate the need to perform validation studies in the populations to which it may be useful to apply these assays, including those where stool sampling is impractical, turnaround times for clinically informative decisions are short, or infrastructure for microbiome sequencing is not available.

Our observations agree with those of published studies, which have found correlations between fecal SBAs and amino acid derivatives with alpha diversity of the gut microbiome.^96–98^ Correlations have also been shown between circulating metabolites and the gut microbiome in other cohorts.^99–104^ For example, Menni and colleagues,^105^ found 3-phenylpropionate (3-PPA), I3P, and cinnamoylglycine all significantly predicted higher diversity, while imidazole propionate predicted lower diversity. Likewise, Diener and colleagues,^102^ found PCS, I3P, hippurate, and all SBA in plasma were significantly associated with gut microbiome composition.

## CONCLUSIONS

Circulating microbial metabolites were consistently associated with gut anaerobe loss and ecological collapse across diverse disease states and changed during treatment-induced gut microbiome disruption, performed comparably to gut microbiome profiling for distinguishing disease from health. While not intended to replace direct gut microbiome assessments using sequencing, our analysis establishes peripheral microbial metabolites as informative biomarkers of gut microbiome disruption. Targeted validation in well-designed observational and interventional cohorts is now required.

## LIST OF ABBREVIATIONS1

3-PPA: 3-phenylpropionate
AML: acute myeloid leukemia
ASCT: allogeneic stem cell transplant
AA: amino acids
Abx: antibiotics
ARVs: anti-retrovirals
AUC: area under the curve
ASVs: Amplicon sequence variants
BIC: Bayesian Information Criterion
CIHR: Canadian Institutes of Health Research
CV: cardiovascular
CENTRAL: Cochrane Central Register of Controlled Trials
CI: confidence interval
DCA: deoxycholate
DC: discharge
DADA2: Divisive Amplicon Denoising Algorithm 2
FDR: false discovery rate
GCDCA: glycochenodeoxycholate
GLCA: glycolithocholate
GUDCA: glycoursodeoxycholate
I3P: indole-3-propionate
IQR: interquartile range
IUDCA: isoursodeoxycholate
LASSO: Least Absolute Shrinkage and Selector Operator
LCA: lithocholate
MetS: Metabolic syndrome
MetaPhlAn: Metagenomic Phylogenetic Analysis
MaAsLin2: Multivariable Association with Linear Models 2
NAFLD: Non-alcoholic fatty liver disease
NA: not applicable
NS: not significant
OR: odds ratio
PAG: phenylacetylglutamine
PCS: p-cresol sulfate
PCG: p-cresol glucuronide
PWH: persons with HIV
PAA: phenylacetate
PEG: polyethylene glycol
PBA: primary bile acids
PCA: principal component analysis
PCoA: principal coordinate analysis
PRISMA: Preferred Reporting Items for Systematic Reviews and Meta-Analyses
QIIME 2: Quantitative Insights Into Microbial Ecology 2
RA: relative abundance
RCT: randomized controlled trials
ROC: receiver operating characteristic
rRNA: ribosomal ribonucleic acid
SBA: secondary bile acids
SRA: sequence read archive
SDI: Shannon diversity index
TMAO: trimethylamine-N-oxide
UDCA: ursodeoxycholate

## DECLARATIONS

### Ethics approval and consent for participation

Not applicable

### Consent for publication

Not applicable

### Availability of data and materials

Compiled microbiome and metabolite data from each study is available as a supplement. The original and full data with accession numbers for extraction from each study is available from the original study and as referenced in the table below. Code generated during this study is available at https://github.com/taylorkain1/metabolome-SR. This study did not generate any new materials or unique reagents.

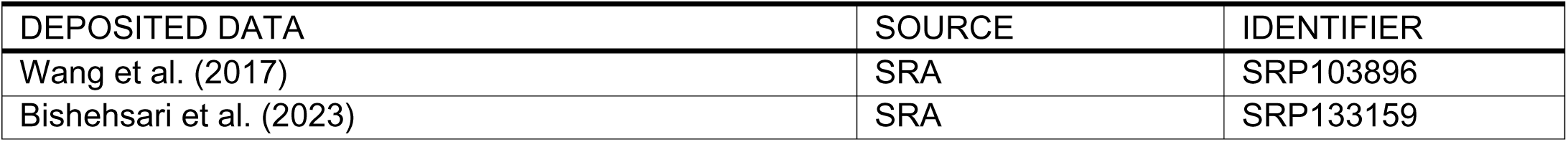
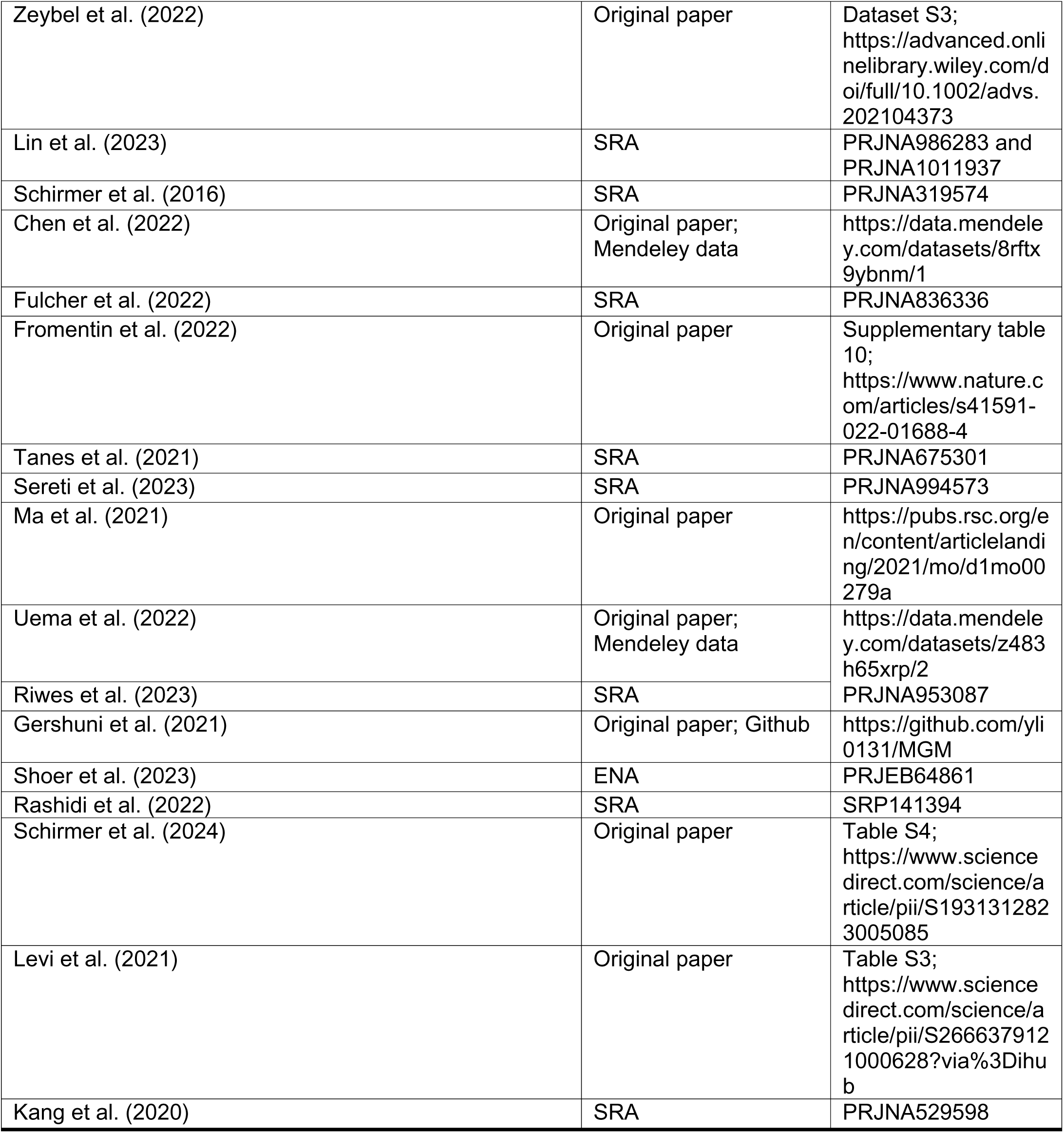

### Author contributions

T.K., and B.C. designed the study; T.K., and E.A. performed the research and analyzed the data. T.K., and E.A. wrote the manuscript; T.K. prepared manuscript for submission.

## Acknowledgements

The authors would like to acknowledge contributions regarding figure generation from other lab members including Noelle Yee, and Maria Kulikova, as well as Dylan Kain for his review and feedback on the completed manuscript. They would also like to acknowledge graduate research support from Canadian Institutes of Health Research (CIHR) and The Department of Medicine at the University of Toronto.

## Authors information

Further information and any requests for primary data should be directed to Bryan Coburn.

## Supplemental Table Titles + Legends for Excel Files

**Supplement Excel File #1 – Box B Studies – Master File.** All extracted metabolite, gut microbiome, and meta-data for all studies used in primary analysis (Box B studies).

**Supplement Excel File #2 – Box A Studies – Overview.** Overview of all papers used in the generation of candidate metabolites with information about size of study, main study population, and links to the original article.

